# Maturing giant kelp develop depth-specific microbiomes

**DOI:** 10.1101/2024.04.04.588117

**Authors:** Sevan Esaian, An Bui, Bartholomew P. DiFiore, Joseph R. Peters, Michelle Lepori-Bui, Kelsey Husted, Holly V. Moeller, Elizabeth G. Wilbanks

## Abstract

Giant kelp (*Macrocystis pyrifera*) is a photosynthetic macroalga that produces dissolved organic carbon (DOC), essential for marine bacteria and food webs. The bacterial communities residing on giant kelp blades consume and compete for complex carbohydrates, contributing to the microbiome community structure. In this study, we investigate how the microbiome changes in response to the age and depth of giant kelp blades and assess how these changes relate to differences in the host’s photophysiology. We find that the microbial community increases in richness and evenness as kelp blades age. While the microbiomes of juvenile blades are stochastic, communities on mature blades coalesce into less variable, depth-specific community types. Differentially abundant genera in mature microbiomes include members of *Bacteroidia* and *Gammaproteobacteria*, known for carbohydrate degradation, and *Planctomycetes*, which often produce protective secondary metabolites. These shifts in microbiome communities are associated with increased maximum quantum yield of photosystem II of mature blades; therefore, they may be linked to enhanced DOC exudation. By shedding light on these dynamics, our study contributes to a better understanding of the complex interplay between macroalgae, their respective microbiomes, and the surrounding marine environment.

## Introduction

Macroalgae play a crucial role in marine ecosystems as the foundation of marine food webs, covering approximately 3.4 million km^2^ of global seabed (Wada *et al*., 2007; Lønborg *et al*., 2009). Beyond their well-studied role as habitat and food sources for marine animals, marine macroalgae make major contributions to the microbial loop, exuding an estimated 1.5 petagrams of carbon per year as dissolved organic carbon (DOC) (Krause-Jensen *et al*., 2018; Sala *et al*., 2019; Chen *et al*., 2020). This DOC is a substantial but highly variable fraction of macroalgal net primary productivity (∼10-60%) and can be an important source in coastal waters (Abdullah and Fredriksen, 2004; Halewood *et al*., 2012; Wada and Hama, 2013). Macroalgal DOC fuels the growth of heterotrophic marine bacteria; however, the fate of this carbon is poorly understood (Lønborg *et al*., 2020; Hall *et al*., 2022).

The largest marine macroalgae, giant kelp (*Macrocystis pyrifera*), exudes ∼14% of its annual net primary productivity as DOC (Dayton, 1985; Reed *et al*., 2015; Krumhansl *et al*., 2016) and supports abundant marine bacteria, both free-living and host-associated (Lin *et al*., 2018; Minich *et al*., 2018; Weigel and Pfister, 2019; James *et al*., 2020). As an abundant carbohydrate source, giant kelp blades foster diverse bacterial heterotrophs containing dozens of bacterial phyla (Lin *et al*., 2018; Minich *et al*., 2018; Weigel and Pfister, 2019; James *et al*., 2020), with population densities approaching 20 million cells per cm^2^ (Tabita Ramírez-Puebla *et al*., 2021). These bacteria colonizing the surface kelp blade surface may play important roles in both the host health and the remineralization of organic carbon.

Prior work has found that the microbial communities associated with the giant kelp canopy vary between geographic sites (Weigel and Pfister, 2019; James *et al*., 2020) and as a function of their host’s physiological condition (James *et al*., 2020), as has been seen in other foundational species of macroalgae (Marzinelli *et al*., 2015; Phelps *et al*., 2021; Wood *et al*., 2022). Studies on different species of kelp and macroalgae demonstrated that both seasonality and hosts anatomy also impact microbiome development(Lemay *et al*., 2021; Davis *et al*., 2023). In mesocosm experiments simulating ocean warming and/or acidification, canopy forming kelp species (*Macrocystis, Ecklonia*) experienced considerable dysbiosis, showing dramatic changes in microbial community composition correlated with tissue damage or decreases in growth (Minich *et al*., 2018). However, the experimental challenges of working with giant kelp have made it difficult to glean mechanistic insight into the origins and consequences of this microbiome variability.

As a canopy-forming species, giant kelp (and its associated microbes) experience dramatic differences in light and temperature as it grows from the seafloor to the water’s surface (Gerard, 1984, 1986). Though the influence of blade age or depth on the kelp microbiome has not yet been described in *Macrocystis*, substantial differences in host physiology with age and depth indicate these factors likely play an important role in microbiome development. For example, the differences in kelp’s photosynthetic capacity and maximum quantum yield lead to considerably higher photosynthetic efficiency and growth rates in surface blades and older blades (Hepburn *et al*., 2007; Edwards and Kim, 2010). Depth can also impact a blade’s ability to exude DOC (Miller *et al*., 2011; Reed *et al*., 2015), which could impact microbial community dynamics.

In photosynthetic organisms, both marine and terrestrial, host aging was marked by a transition from highly variable microbiomes amongst juveniles to mature microbiomes that both richer, more even and less variable and found greater compositional stochasticity in juvenile-associated microbiomes (Wagner *et al*., 2016; Sanders-Smith *et al*., 2020). Similarly, studies of kelps where annual blades grow continuously (e.g. *Nereocystis* and *Laminaria* species) have found that older tissue host richer bacterial communities than newly synthesized meristematic tissues (Bengtsson *et al*., 2011; Weigel and Pfister, 2019; Lemay *et al*., 2021). In *Macrocystis pyrifera*, blades grow to a maximum length of 80 centimeters (Abott and Hollenberg, 1976)and have typical lifespans ranging from 40 to 90 days (Rodriguez *et al*., 2016). We hypothesize that blade age and its depth environment have potential synergistic effects that are likely to influence the patterns of microbial community assembly.

Here, we demonstrate that mature giant kelp blades have greater photosynthetic efficiency and capacity than their juvenile counterparts at all depths. These photophysiological changes are correlated with an increase in the richness of the microbial communities associated with mature blades, which unlike their juvenile counterparts, coalesce into depth-specific microbiome-types. We find this development of depth-specific mature microbiomes is driven by a small subset of genera.

## Methods

### Sampling

We collected giant kelp blades in June and July 2019 from Arroyo Quemado reef, a long-term ecological research (LTER) site located in the Santa Barbara channel (34°28’07.6”N 120°07’17.1”W). Through the sampling campaign, *in situ* sensors recorded an average sea surface temperature of 21°C and a benthic temperature of 12°C. We concentrated our sampling within a 15 m horizontal radius at the center of the reef (34°28’07.6”N 120°07’17.1”W) to minimize geospatial variation in giant kelp microbiome community composition.

To generate a depth-stratified set of samples, we categorized giant kelp blades into three groups: surface (floating on the water surface), middle (2 - 7 m depths), and bottom (>7m depth). To follow the development of giant kelp blades by age, we tagged 30 fronds per depth category, positioning the tags 10 juvenile blades (i.e., scimitars) back from the growing tip, and then destructively sampled blades from a subset of these marked fronds at each time point. After 2 weeks, we collected a total of 45 newly grown, two-week old blades (3 blades per frond and 5 fronds per depth category). We repeated the same procedure 2 weeks later to obtain 4-week-old blades. During collections, we placed the giant kelp blades in Whirl-Pak bags (Fisher Scientific) filled the bags with seawater and sealed them while keeping the pneumatocyst outside of the bags.

We sampled microbes from the tip, center, and base of each blade’s surface using a closed-circuit syringe as previously described (Haas *et al*., 2014; Lim *et al*., 2014; James *et al*., 2020). We avoided regions with epiphytes or their calcified remnants, as these regions are known host different microbial communities (James *et al*., 2020). For each blade, we filtered a total of 150 mL of syringe volume through a single 0.2 µm polyethersulfone filter cartridge (Sterivex-GP, Millipore). We then filled each cartridge with 1 mL of sucrose lysis buffer and stored them at - 20°C until DNA extraction.

To determine microbial composition in the environment, we collected 2 L of seawater alongside sampled giant kelp blades. For each depth category on each sampling day, we filtered 500 mL of whole seawater onto 0.2 µm polyethersulfone filter cartridges (Sterivex-GP, Millipore). After filtering, we filled each cartridge with 1 mL of sucrose lysis buffer and stored them at -20°C until DNA extraction.

### Quantifying giant kelp blade surface area

To determine the surface area of giant kelp blades, we captured images of both sides of each blade against a gridded background. Using ImageJ (Fiji) we manipulated the color threshold of each image, removing the background and isolating the blades. We calculate the average surface area (cm^2^) of each blade using images from both sides, resulting in a single value per blade for further analysis.

### Quantifying giant kelp blade maximum quantum yield of photosystem II (Fv/Fm)

To determine the maximum quantum yield of photosystem II, we dark-acclimated all blades for 15 minutes in the laboratory, after microbiome sampling. Using a junior-PAM (Walz), we measured the *Fv/Fm* of each blade. Measurements were taken in 1-inch intervals from the pneutmatocyst to the blade tip, with triplicate readings at each location. We calculated the average of these *Fv/Fm* readings from each blade, resulting in a single value per blade for further analysis.

### Chlorophyll extraction and pigment measurement

In the laboratory, we excised 3 tissue samples from each blade’s tip, center, and base and extracted chlorophyll as described previously (Seely *et al*., 1972; Bell *et al*., 2018). Briefly, cells were lysed in DMSO, rinsed with water, followed by a final extraction in acetone, methanol, and water. We used a fluorometer to measure chlorophyll concentration and calculated the total chlorophyll per unit area for each sample. Each blade resulted in one Chl*a*:C measurement used in subsequent analyses.

### Giant kelp blade carbon and nitrogen measurements

To quantify the percentage of carbon and nitrogen per blade, we excised and combined 5 cm^2^ punches from the base, center, and tip of each blade. We sent tissue samples to Brookside Laboratories, where carbon and nitrogen content were measured using an EL cube elemental analyzer. Each blade resulted in one C:N value used in subsequent analyses.

### DNA extraction and 16S rRNA sequencing

DNA was extracted from filter cartridges as described previously (Wear *et al*., 2018; James *et al*., 2020). Briefly, filters were thawed on ice, cells were lysed using 10% SDS and proteinase K (20 mg/mL), and DNA was extracted using phenol:chloroform:isoamyl alcohol (25:24:1, Thermo Fisher Scientific). DNA was ethanol precipitated and fluorometrically quantified (Qubit, Thermo Fisher Scientific). 16S rRNA genes were amplified using the 515F (GTGYCAGCMGCCCGCGGTAA) and 806R-B (GGACTACNVGGGTWTCTAAT) primers with one-step PCR to generate bacterial 16S V4 amplicons (Wear *et al*., 2018). The resulting amplicons were gel purified (QIAquick Gel Extraction Kit, Qiagen) and normalized (SeQualPrep normalization plate kit, ThermoFisher Scientific) before sequencing the barcoded amplicons on the Illumina MiSeq platform with 300 bp paired end (PE) reads at the University of California, Santa Barbara California NanoSystems Institute.

### 16S rRNA pipeline and analysis set-up

We processed sequence reads using the DADA2 pipeline in R (Callahan *et al*., 2016). Forward reads were trimmed to 200 bp, and reverse reads were trimmed to 160 bp based on sequence quality. Taxonomy was assigned to amplicon sequence variants (ASVs) using the SILVA taxonomy database (v.132). We excluded ASVs identified as mitochondria, chloroplasts, and eukaryotes (Quast *et al*., 2013). After this quality filtering the sequence reads, we obtained a total of 10^7^ reads from 90 samples, resulting in 7,600 distinct ASVs. The sequence read counts per sample ranged from 26,000 to 200,000. Due to this substantial variation in read counts, we rarefied the ASV counts for further analysis using DADA2 (Callahan *et al*., 2016).

### Analyzing differences in blade photophysiology and microbiome composition by age and depth

Sequence and photophysiology data were processed and plotted in R using the *tidyverse* package (Wickham *et al*., 2019). To assess potential significant variations in giant kelp blade photophysiology based on age and depth categories, we used the *vegan* R package (Oksanen, 2022) to conduct an ANOVA and Tukey’s HSD post-hoc tests. In this analysis, age and depth categories served as independent variables, while each aspect of photophysiology was considered a dependent variable.

After rarefaction, we computed the percent relative abundance of each ASV in each sample. These values were utilized for the calculation of diversity metrics and subsequent multivariate statistical analyses using *vegan*. Our analysis involved ASV richness, Pielou’s Evenness, Shannon Diversity, Simpson Diversity, and beta dispersion (indicating community turnover across blades).

To further explore these compositional differences, we employed constrained (Unifrac weighted) principal component analysis (PCA), canonical correspondence analysis (CCA), non-scaled principal coordinate analysis (PCoA), and unconstrained non-metric multidimensional scaling (NMDS, Bray-Curtis) using *vegan*. We employed these different ordinations to identify robust patterns that were consistent across methodological approaches. We calculated significant differences in age and depth categories from NMDS outputs using PERMANOVA, ANOSIM, and beta dispersion. We utilized PERMANOVA further to quantify significant patterns in photophysiology characteristics combined with age and depth categories based on microbiome NMDS coordinates.

### Modeling drivers of microbiome shifts across age and depth categories

To characterize changes in the microbiome across age and depth, we employed the R packages *phyloseq* (McMurdie and Holmes, 2013) to create phyloseq objects and *corncob* (differentialTest and bbdml) to calculate differential abundance using abeta-binomial regression at the genus- and ASV level (Martin *et al*., 2020). To compare juvenile and mature samples, a model was constructed to test differential abundance and variability between age categories while controlling for the effect of depth on dispersion (formula=depth+age; *phi.formula*=depth; *formula_null*=age; *phi.formula_null*=1).

In a separate analysis, we investigated the effect of depth on changes in taxon abundance, and we focused solely on mature blade microbiomes because juvenile samples did not exhibit significant differences in community structure based on depth (Supp. Table 1). As the model designed to handle binary comparisons (*e.g.* surface vs. subsurface), we combined samples from middle and bottom depth into a subsurface category, which significantly differed from mature surface samples (Supp. Table 1). Here, our model tested for differential abundance and variability between depth categories, while accounting for overdispersion (*formula*=depth; *phi.formula*=depth; *formula_null*=1; *phi.formula_null*=1). After quantifying relative abundances using beta-binomial regression, we filtered out genera with less than 0.5% relative abundance. From each model, we retained genera that exceeded this cutoff and proceeded to model the differential abundance of their corresponding ASVs. We applied a prevalence cutoff where the ASV of interest must have a relative abundance greater than 0 in at least five samples. Then, we examined the differential abundance of each ASV across age or depth categories (Tables 1, Supp. Fig. 5).

**Table 1:**
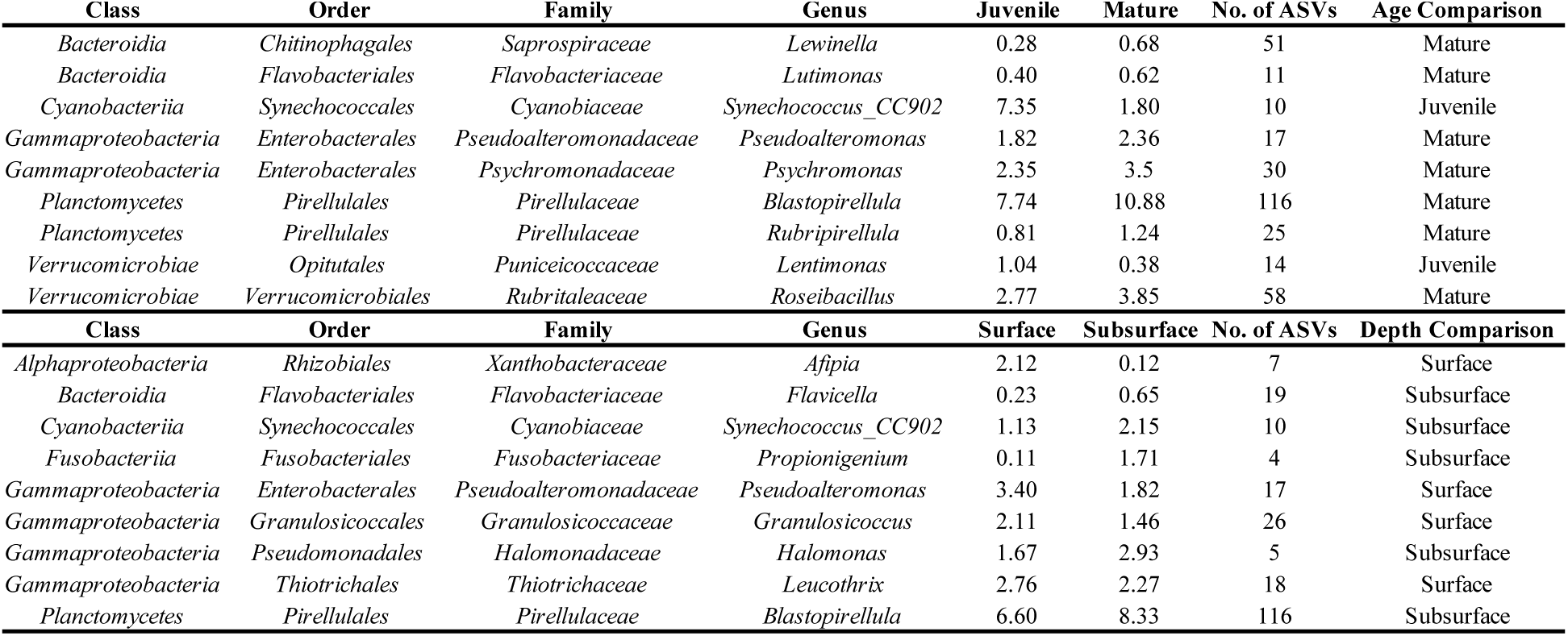
Several bacterial genera drive differences in giant kelp microbiome composition as a function of blade age and depth of mature samples. Genera shown are those that had a significant differential abundance by either age (top) or depth (bottom), as identified by a beta binomial regression (*P*<0.05). Columns display the statistically significant genera, their average percent relative abundance, the number of ASVs in each genus, and the category with significantly greater percent relative abundance.

## Results

### Giant kelp blade photophysiology changes by age and depth

The surface area of giant kelp blades was similar across categories, except for juvenile blades at middle depths which were significantly larger (Fig. 1A). Mature blades had significantly higher maximum quantum yield of photosystem II and higher Chl*a*:C ratio compared to their juvenile counterparts (Fig. 1B and 1C). For juvenile blades the maximum quantum yield decreased with depth, unlike mature blades which had consistent and maximum high quantum yields across all depth categories (Fig. 1B). We found that surface blades had enriched in C:N ratios compared to samples from deeper depths, but there were no significant differences between age categories (Fig. 1D).

**FIGURE 1.**
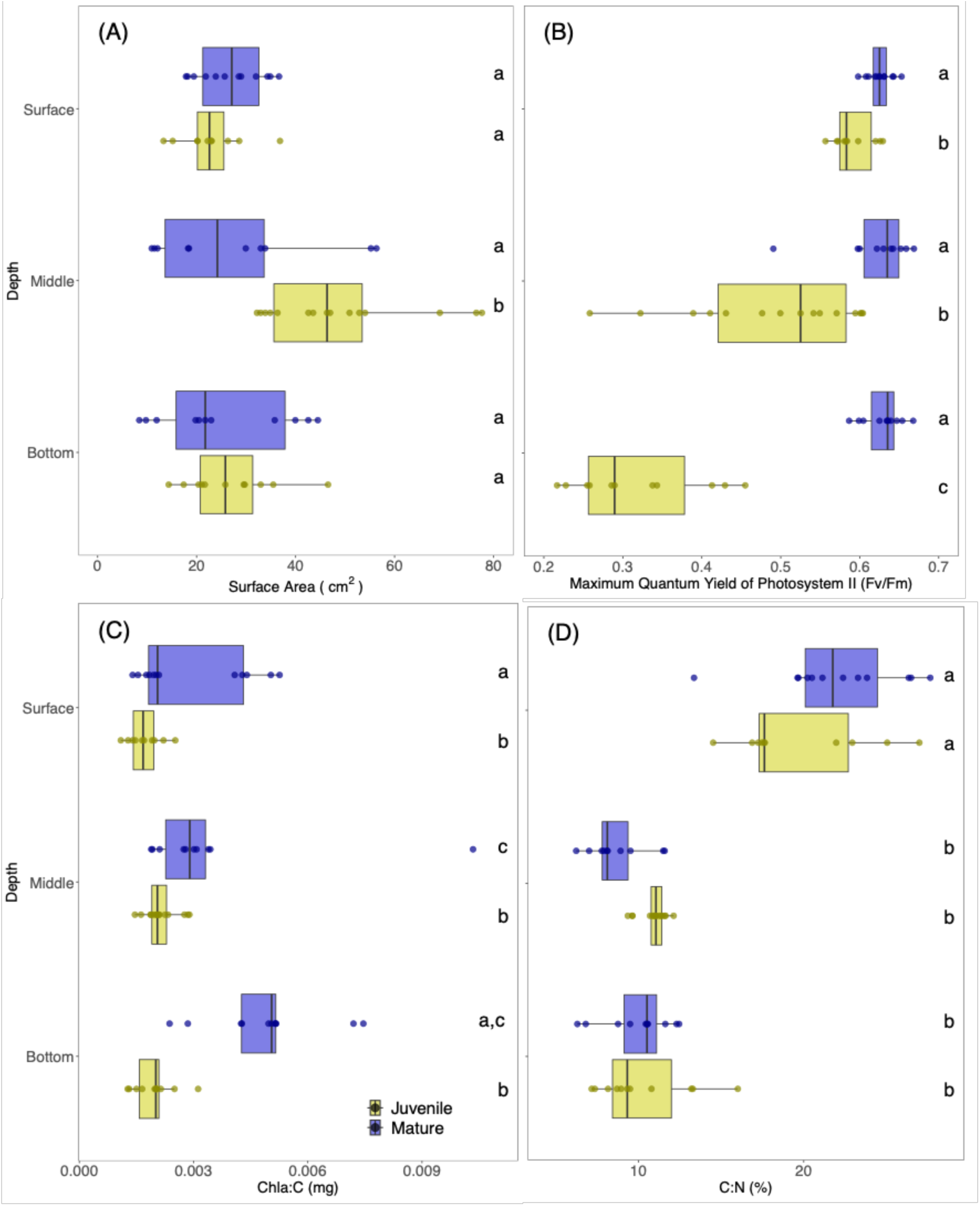
Giant kelp blades have photophysiological differences as a function of both age and depth. Boxplots depict the spread of measurements for independent photophysiological characteristics in giant kelp blades across age (juvenile shown in yellow and mature in blue) and depth categories. Points represent one measurement per blade and solid black line represents median. Significant differences, denoted by letters at right, were determined using ANOVA and Tukey’s HSD post-hoc test (*P*<0.05).

### Microbiome compositional changes by age and depth

Across multiple ordination methods, juvenile microbiomes were quite stochastic (Fig. 2, Supp. Fig. 2) with greater dispersion than mature blades (Fig. 3C). Juveniles exhibited no significant differences in community composition across depths (Supp. Table 1). In contrast, mature blade microbiomes converged onto significant depth-specific compositions (Fig. 2, Supp. Table 1) that were consistent across multiple ordination methods (Supp. Fig. 2). Across all samples, the giant kelp microbiome was significantly different from the free-living communities which were quite similar through depths and time (Supp. Fig. 3 and Supp. Table 1).

**FIGURE 2.**
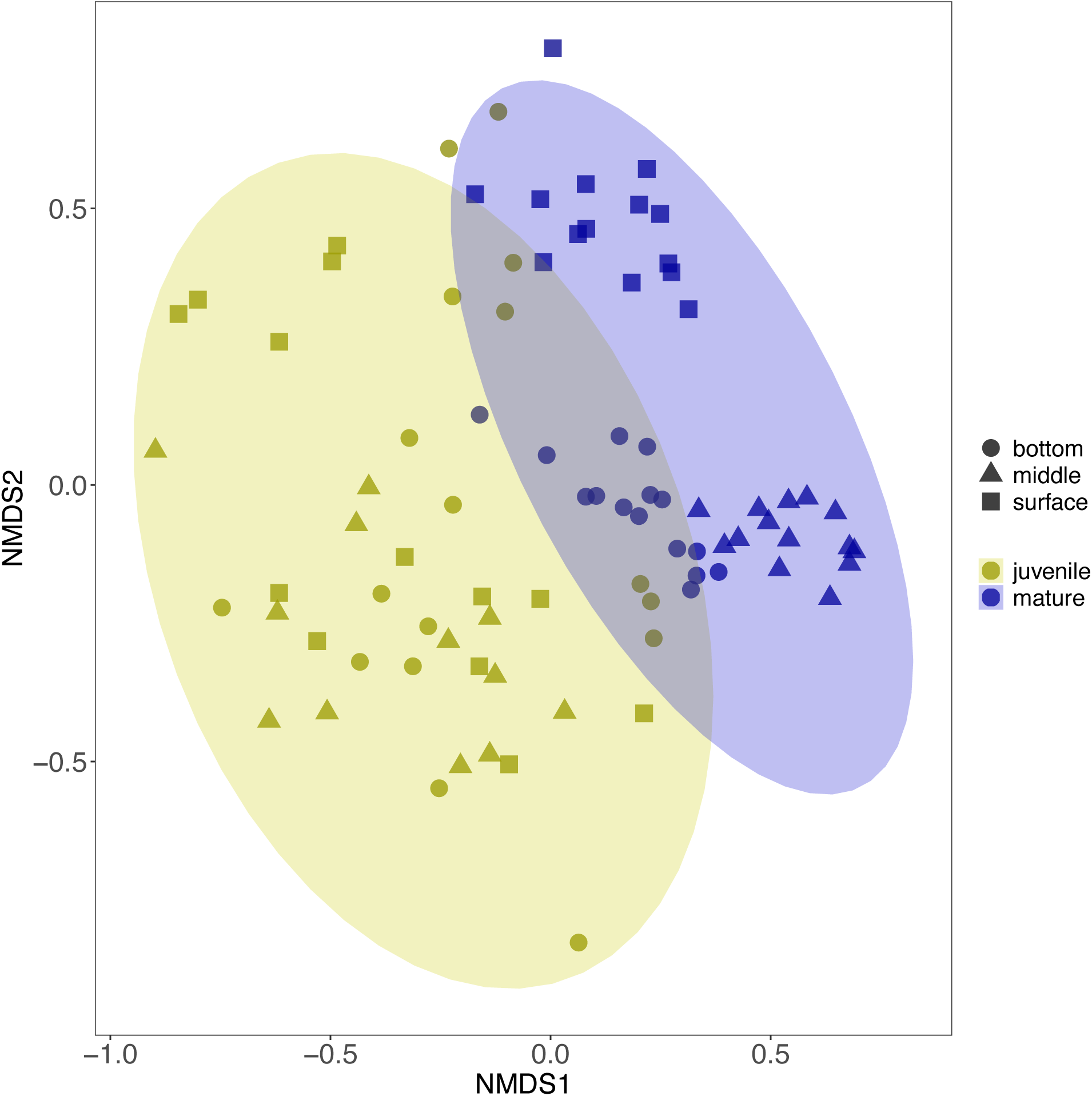
Giant kelp develops depth specific microbial communities as blades age. Non-metric multidimensional scaling (NMDS) model of percent relative abundances microbial communities from juvenile and mature giant kelp blades (yellow and blue, respectively) sampled from three depths (circle, triange, squares). The analysis employs Bray-Crutis distances, with ellipse drawn at 0.95 cutoff.

**FIGURE 3.**
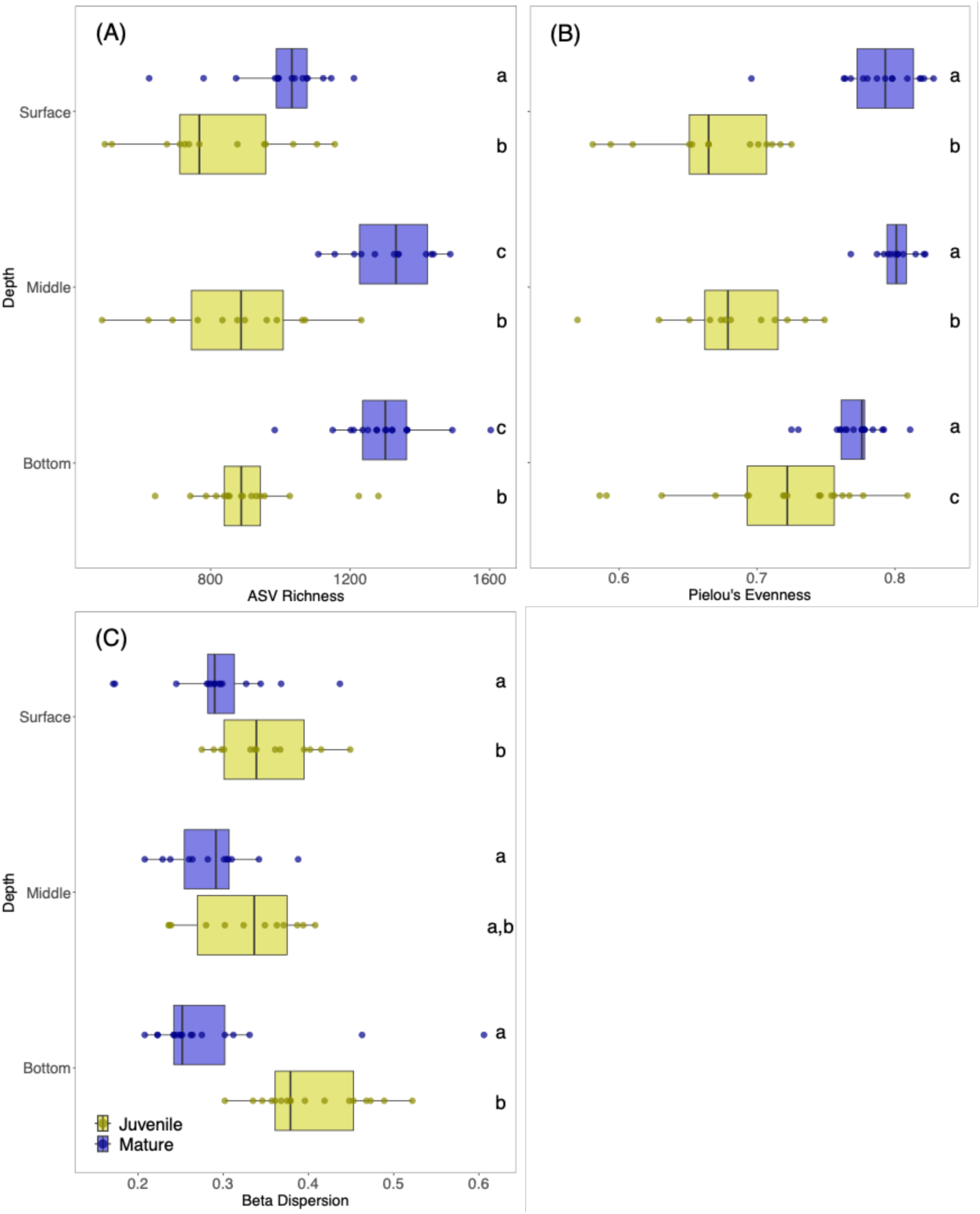
Microbial communities become richer, more even, and less dispersed as giant kelp blades age. Boxplots depicting spread of ASV richness (A), Pielou’s Evenness (B), and beta dispersion (C) in microbial communities from juvenile and mature giant kelp blades (yellow and blue, respectively) sampled from three depths. Points represent one value per microbiome and solid black line indicates median for that dataset. Significant differences are denoted by letters determined using ANOVA and Tukey’s HSD post-hoc test (*P*<0.05).

As giant kelp blades aged, there was a significant increase in ASV richness and Pielou’s Evenness, accompanied by a decrease in beta dispersion in the microbiomes (Fig. 3). Mature middle and bottom blades exhibited the highest ASV richness (Fig. 3A). Through CCA, robust relationships emerged between the microbiome community compositions of mature blades and photophysiology characteristics including maximum quantum yield of photosystem II (*Fv/Fm*) and Chl*a*:C (Fig. 4). Similarly, significant associations were observed between the microbiomes of surface blades and the C:N ratio (Fig. 4). We also found that maximum quantum yield of photosystem II (*Fv/Fm*) in combination with either age or depth was a significant driver of microbiome composition (Supp. Table 2).

**FIGURE 4.**
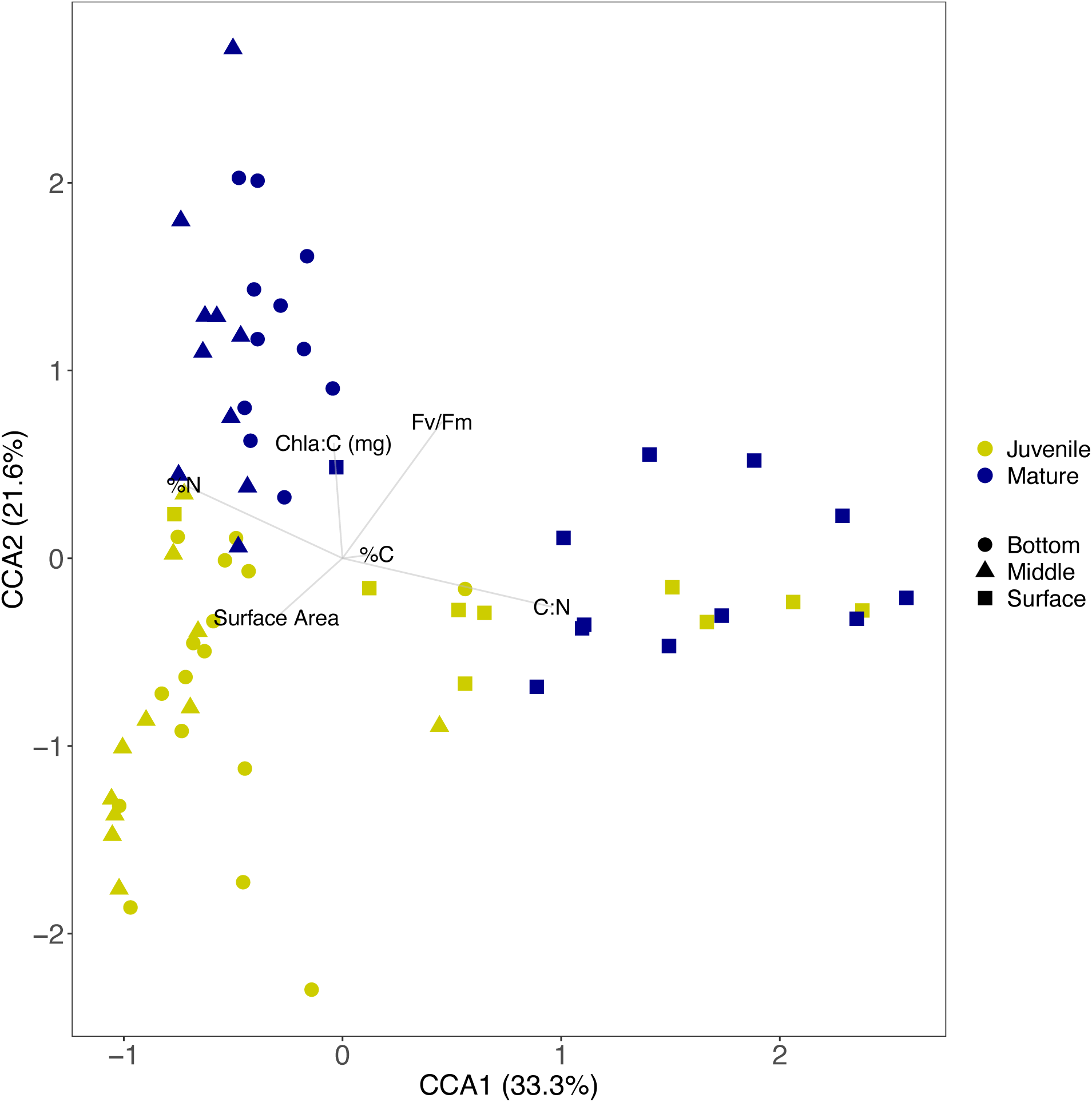
Photophysiological metrics are associated with the development of depth specific microbiomes on mature giant kelp blades. Canonical Correspondence Analysis (CCA) model of percent relative abundance of giant kelp blade microbiomes and the respective photophysiological characteristics of each blade. Samples include juvenile and mature giant kelp blades (yellow and blue, respectively) sampled from three depths (shapes). Arrow length indicates strength and direction of the relationship, while the axes represent the proportion of variation explained.

### Dominant bacterial genera and ASVs shaping microbiome composition

In total, there were 15 bacterial orders with average relative abundances greater than 1% across all samples (Supp. Fig. 6). These include taxa commonly associated with macroalgal surfaces and reported in prior surveys of the giant kelp microbiome, such as members of the *Planctomycetes, Verucomicrobiales, Caulobacterales, Rhodobacterales* and *Bacteroidia* (Lachnit *et al*., 2011; Lage and Bondoso, 2014; Weigel and Pfister, 2019; Tabita Ramírez-Puebla *et al*., 2021). We detected 26 bacterial genera spanning 6 phyla, that had an average relative abundance of at least 0.5% across all samples (Supp. Table 3). To elucidate drivers of the statistically significant community composition differences based on age alone (Fig. 2, Supp. Fig. 2, and Supp. Table 1), we pooled samples across depths and identified genera that were differentially abundant between juvenile and mature samples. We found 9 genera that exhibited statistically significant differences in their abundance across age categories (Table 1 and Supp. Fig. 4A). Genera with higher relative abundances in juvenile samples included *Synechococcus_CC9902* and *Lentimonas*, while those with greater relative abundance in mature samples included *Lewinella*, *Lutimonas*, *Pseudoalteromonas*, *Psychromonas*, *Blastopirellula*, *Rubripirellula*, and *Roseibacillus* (Table 1 and Supp. Fig. 4A). All genera belonged to a dominant bacterial order (Supp. Fig. 6). *Lentimonas* and *Psychromonas* exhibited differences driven by only two ASVs, whereas *Blastopirellula* contained five ASVs with increased abundances in mature microbiomes (Supp. Fig 5A and Supp. Table 4).

To examine depth-related shifts in microbiome composition, we focused our analysis on mature samples due to the stochastic nature of juveniles (Supp. Table 1). We categorized blades into surface and subsurface groups because their compositional patterns were significantly different and beta-binomial regressions permit two categories (Fig. 2 and Supp. Table 1). Beta-binomial regression analyses revealed higher relative abundances of *Afipia*, *Leucothrix*, *Pseudoalteromonas*, and *Granulosicoccus* at the surface, while *Flavicella*, *Synechococcus_CC9902*, *Propionigenium*, *Halomonas*, and *Blastopirellula* showed higher abundances in subsurface microbiomes (Table 1, Supp. Fig. 4B and 5B, and Supp. Table 4). With the exception of *Propionigenium*, all genera belonged to a dominant bacterial order (Supp. Fig. 6). For both age and depth comparisons of the *Blastopirellula*, we found that while genus-level trends were supported by most ASVs (e.g. higher relative abundance at depth or in mature blades), we also identified some statistically significant ASVs that were opposite that of the majority (e.g. higher relative abundance at the surface or in juvenile blades). This nuanced ASV-level variation within a genus underscores the complexity of microbiome dynamics and highlights the importance of considering individual ASVs when interpreting compositional changes.

## Discussion

The first portion of this study aimed to assess giant kelp blade dynamics across depths and age, focusing on photophysiological characteristics. Mature blades exhibited higher maximum quantum yields of photosystem II and Chl*a*:C compared to juveniles, supporting previous findings (Fig. 1) (Edwards and Kim, 2010; Tom W. Bell *et al*., 2015; Bell *et al*., 2018; Weigel *et al*., 2022). Chl*a*:C measurements aligned with prior research, while C:N values fell within reported ranges, suggesting the kelp are absorbing nitrogen from benthic invertebrate excretions (Fig. 1D; Fig. 4) (Tom W. Bell *et al*., 2015; Tom W Bell *et al*., 2015; Bell *et al*., 2018; Peters *et al*., 2019).

The second part of our study analyzed changes in the giant kelp microbiome communities concerning age and depth, revealing pronounced differences, particularly in mature blades (Fig. 2; Supp. Table 1; Supp. Fig 2). Mature giant kelp blade microbiomes exhibited greater similarity within each depth, indicating a shift from stochastic to depth-specific communities as blades aged (Fig. 2A; Fig. 3C; Supp. Fig. 3), emphasizing the considerable influence of local environmental conditions, especially depth, on microbial community assembly (Fig. 4) (Wagner *et al*., 2016; James *et al*., 2020; Sanders-Smith *et al*., 2020). As giant kelp blades mature, the richness and evenness in their microbiomes increases, consistent with findings in other kelps and plants, both marine and terrestrial(Wagner et al., 2016; Sanders-Smith et al., 2020). However unlike our findings on *Macrocystis* and prior work on plants, studies on kelp species characterized by continuous of annual blades (e.g. *Nereocystis* and *Laminaria*) did not observe higher dispersion amongst the microbiomes sampled from juvenile tissues (Bengtsson *et al*., 2011; Weigel and Pfister, 2019; Lemay *et al*., 2021). The mechanism underlying this difference is unclear, but clearly reflects a differences in microbial colonization and community succession patterns of continuously growing host tissue compared to leaves and blades with a limited lifespans and a maximum mature size.

Both age and depth exerted influence on the abundance of bacterial taxa associated with giant kelp blades (Fig. 3). Except for *Synechococcus_CC9902*, all differentially abundant genera identified in our analysis were previously reported as dominant members of giant kelp microbiomes (Lin *et al*., 2018; Minich *et al*., 2018; Weigel and Pfister, 2019; James *et al*., 2020). Mature and subsurface microbiomes showed greater proportions of *Bacteroidia* (Tables 1; Supp. Fig 4) (Lin *et al*., 2018; Weigel *et al*., 2022). Many *Bacteroidia* species utilize high molecular weight organic carbon compounds and possess filamentous cells capable of penetrating host cell walls to access carbohydrates within kelp blade meristoderm (Tabita Ramírez-Puebla *et al*., 2021). Mature blades also had an increase in the relative abundance of gammaproteobacterial genera (*Pseudoaltermonas, Psychromonas*; Table 1), known for their diverse carbohydrate degradation capabilities, consistent with observations from controlled experiments where *Gammaproteobacteria* significantly increased in giant kelp microbiomes under higher temperature and *p*CO_2_ conditions (Minich *et al*., 2018).

In our study, *Planctomycetes* (*Blastopirellula* and *Rubripirellula*) are more abundant in mature microbiomes, driven by several distinct ASVs (Table 1; Supp. Fig 4). *Planctomycetes* are abundant across diverse macroalgal microbiomes, due to their adaptations for surface attachment (holdfasts) and their ability to utilize sulfated macroalgal polysaccharides (e.g. fucan, laminarinan) (Lage and Bondoso, 2014). Their prolific arsenal of bioactive compounds may benefit the host by shaping the microbiome and reducing biofouling of the surface (Lage and Bondoso, 2014; Graça *et al*., 2016). Furthermore, other studies have shown that *Pirellulaceae* species are environmentally resilient and remain abundant in microbiomes from different kelp species and across habitats with variable abiotic conditions (Davis et al., 2023 Add Weigel and Pfister 2019).

Examining shifts in the giant kelp microbiome composition as the host ages is crucial for understanding the host-microbiome relationship and the fate of exuded DOC. Previous studies on multiple macroalgal species have demonstrated significant local effects (Lin *et al*., 2018; Weigel and Pfister, 2019; James *et al*., 2020; Davis *et al*., 2023), and our findings reveal clear depth and age effects. Depth-specific differences in the microbial communities associated with mature blades suggests potential a mechanism for differences DOC remineralization throughout the water column. As host blades senesce, changes in the types of carbohydrates they exude, such as an increase in fucoidan, could likely impact the microbiome composition by favoring carbohydrate specialists (Zhang *et al*., 2024). This research elucidates the assembly of microbial communities on healthy giant kelp, a key step in building a mechanistic understanding of the microbiome’s role in host health and biogeochemical cycling in coastal environments.

## Supporting information

All Supplementary Figures and Tables

## Acknowledgements

We completed this project with support from: NSF Grant OCE-1831937, Schmidt Environmental Solutions Fellowship, Coastal Fund Fall 19-12, UCSB startup funds and a Senate Research Grant to HVM, and UCSB startup funds to EGW. We would like to thank Christoph Pierre (UCSB Marine Operations) for sample collection and Jennifer Smith (UCSB Biological Nanostructures Laboratory) for 16S rRNA gene sequencing. Use was made of computational facilities purchased with funds from the National Science Foundation (CNS-1725797) and administered by the Center for Scientific Computing (CSC). The CSC is supported by the California NanoSystems Institute and the Materials Research Science and Engineering Center (MRSEC; NSF DMR 2308708) at UC Santa Barbara.

## Notes

### Competing Interest Statement

The authors have declared no competing interest.

