## Supplementary material for "Maturing giant kelp develop depth-specific microbiomes": All Supplementary Figures and Tables

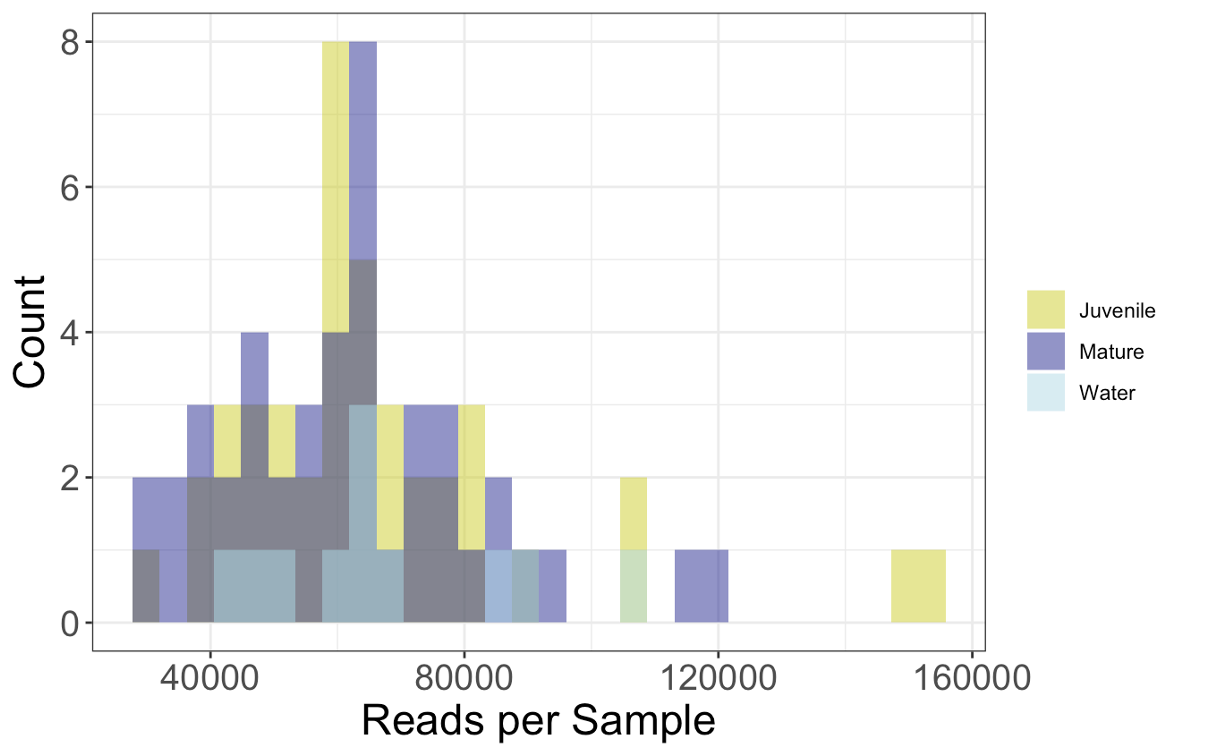

**SUPPLEMENTAL FIGURE 1:** Histogram depicting the distributions of 16S V4-V5 raw sequence reads per sample in juvenile giant kelp blade microbiomes, mature giant kelp blade microbiomes, and free-living seawater bacterial communities. Each category encompasses samples from all three depth bins.

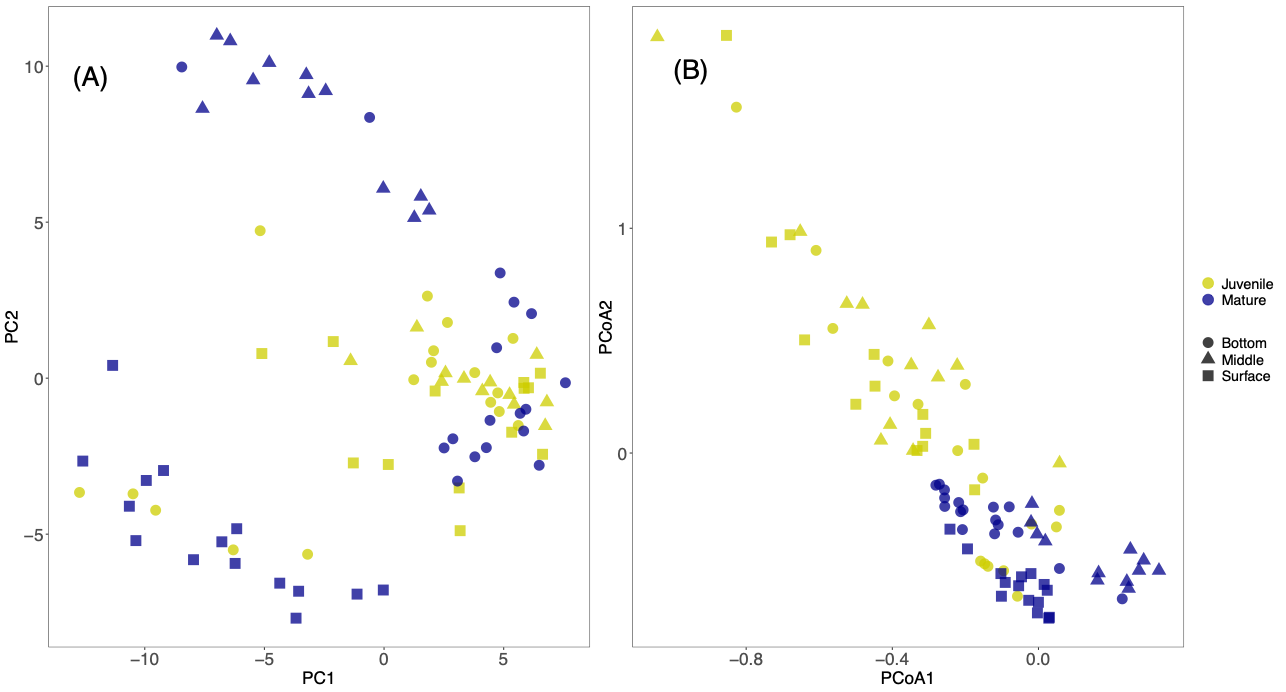

**SUPPLEMENTAL FIGURE 2:** (A) Principal component (not scaled) analysis of percent relative abundances of giant kelp microbiomes across age and depth categories. (B) Principal coordinate analysis (UniFrac weighted) of percent relative abundances of giant kelp microbiomes across age and depth categories.

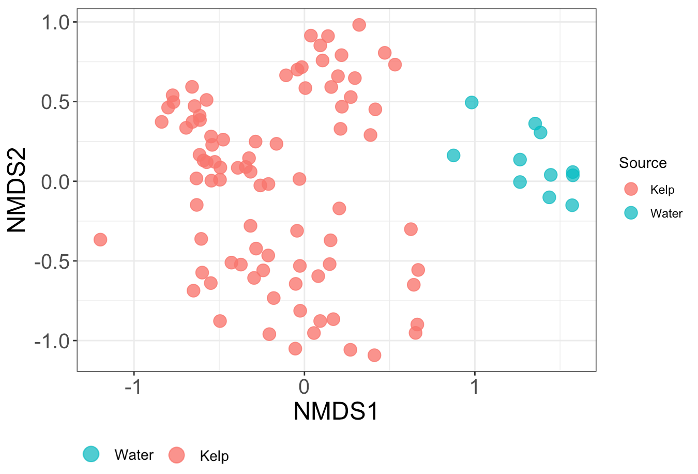

**SUPPLEMENTAL FIGURE 3:** Non-metric multidimensional scaling (NMDS) model of percent relative abundance of giant kelp microbiomes and free-living sea water communities. The analysis employs Bray-Curtis distances.

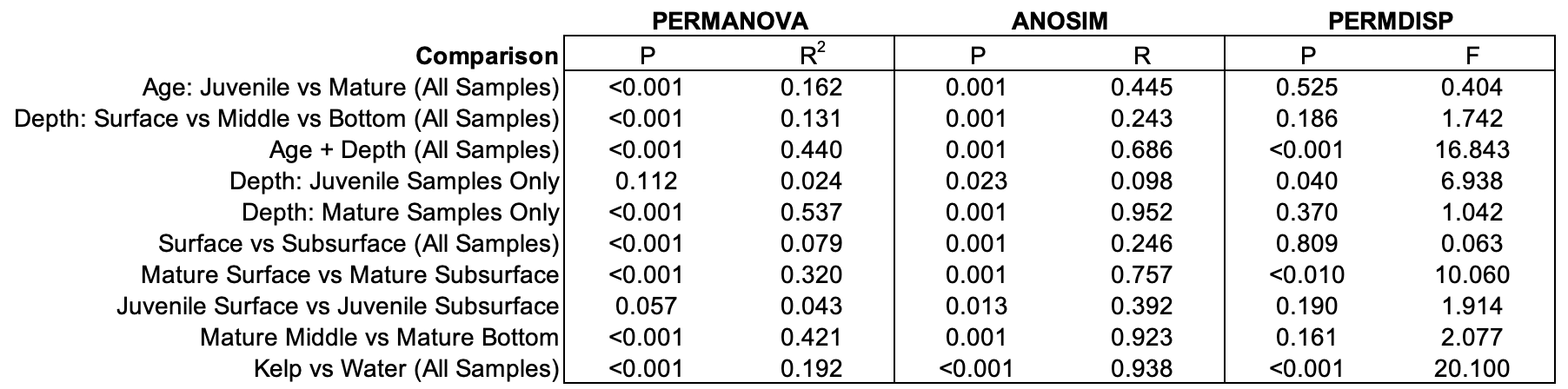

**SUPPLEMENTAL TABLE 1:** Permutational analysis of variance (PERMANOVA), analysis of similarities (ANOSIM), and permutational analysis of multivariate dispersions (PERMDISP) analyses based on percent relative abundances of giant kelp microbiomes. Analyses are separated by comparison type.

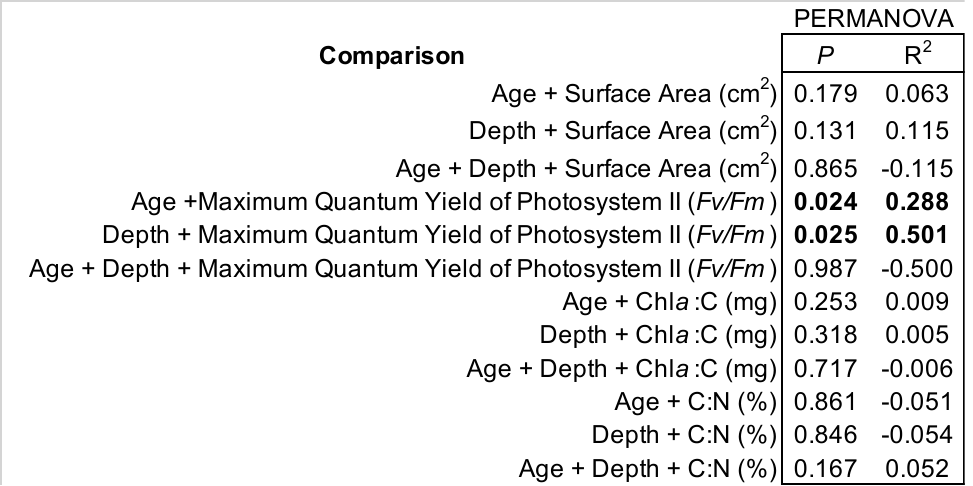

**SUPPLEMENTAL TABLE 2:** Permutational analysis of variance (PERMANOVA), based on giant kelp blade age, depth, photophysiology measurements, and microbiome NMDS coordinates based on percent relative abundance. Bold indicates statistically significant differences.

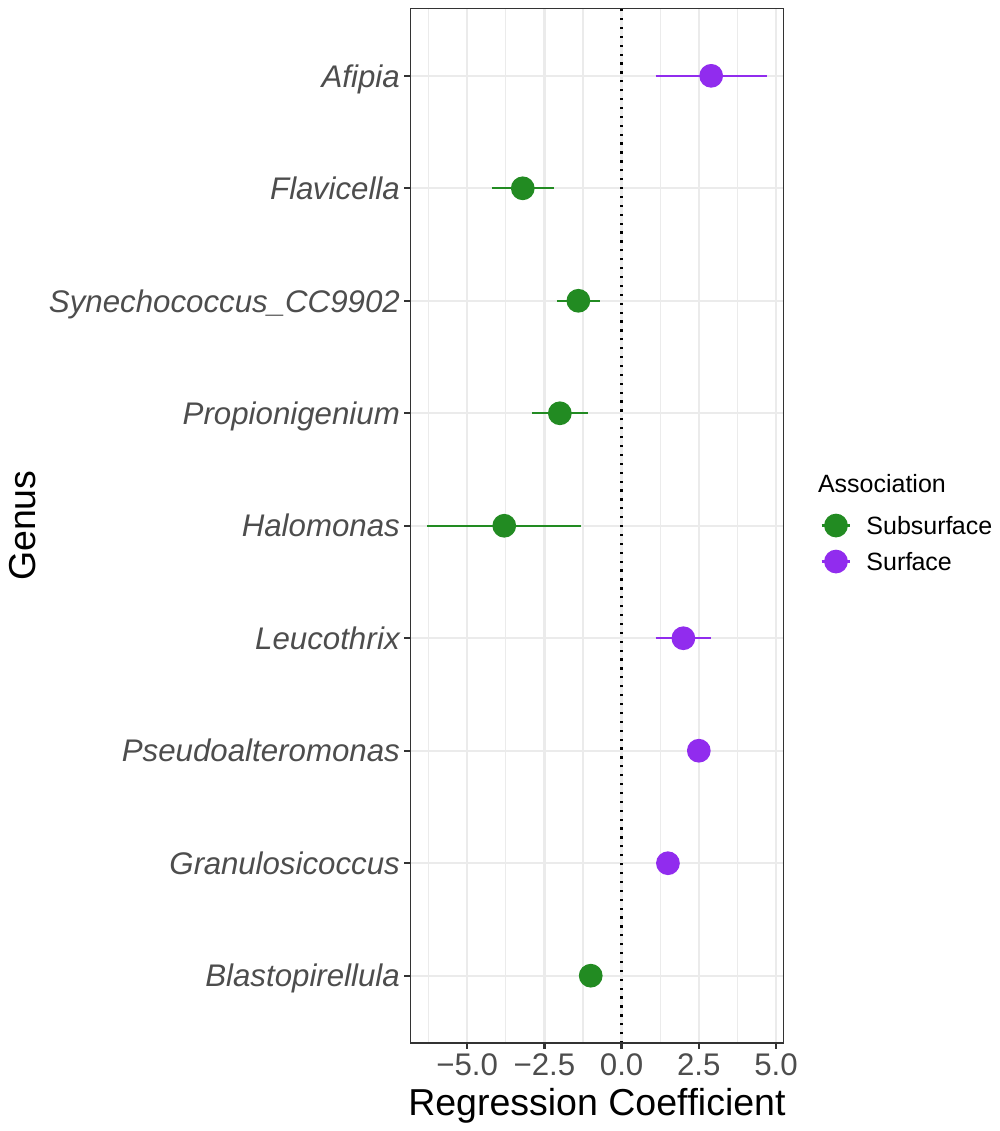

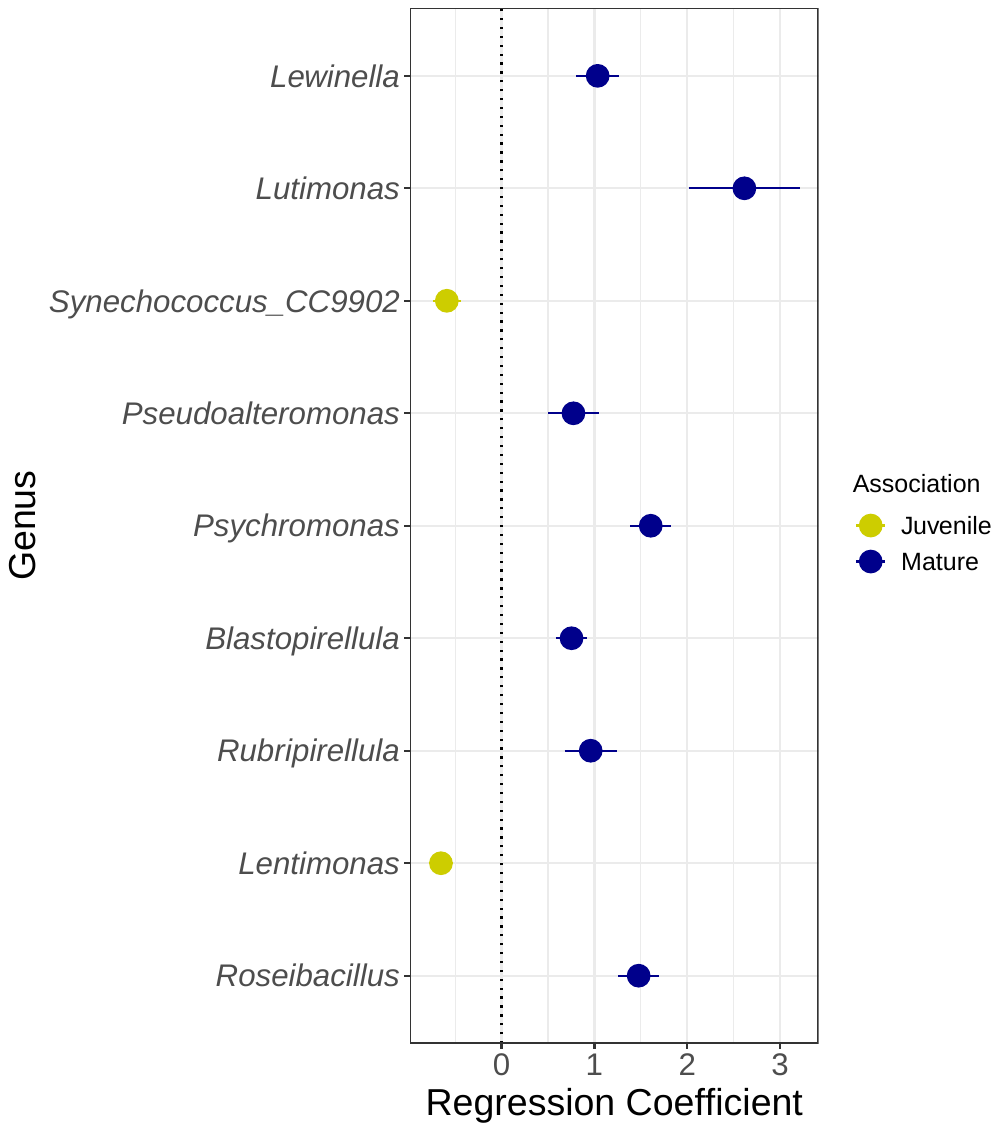

A

B

A

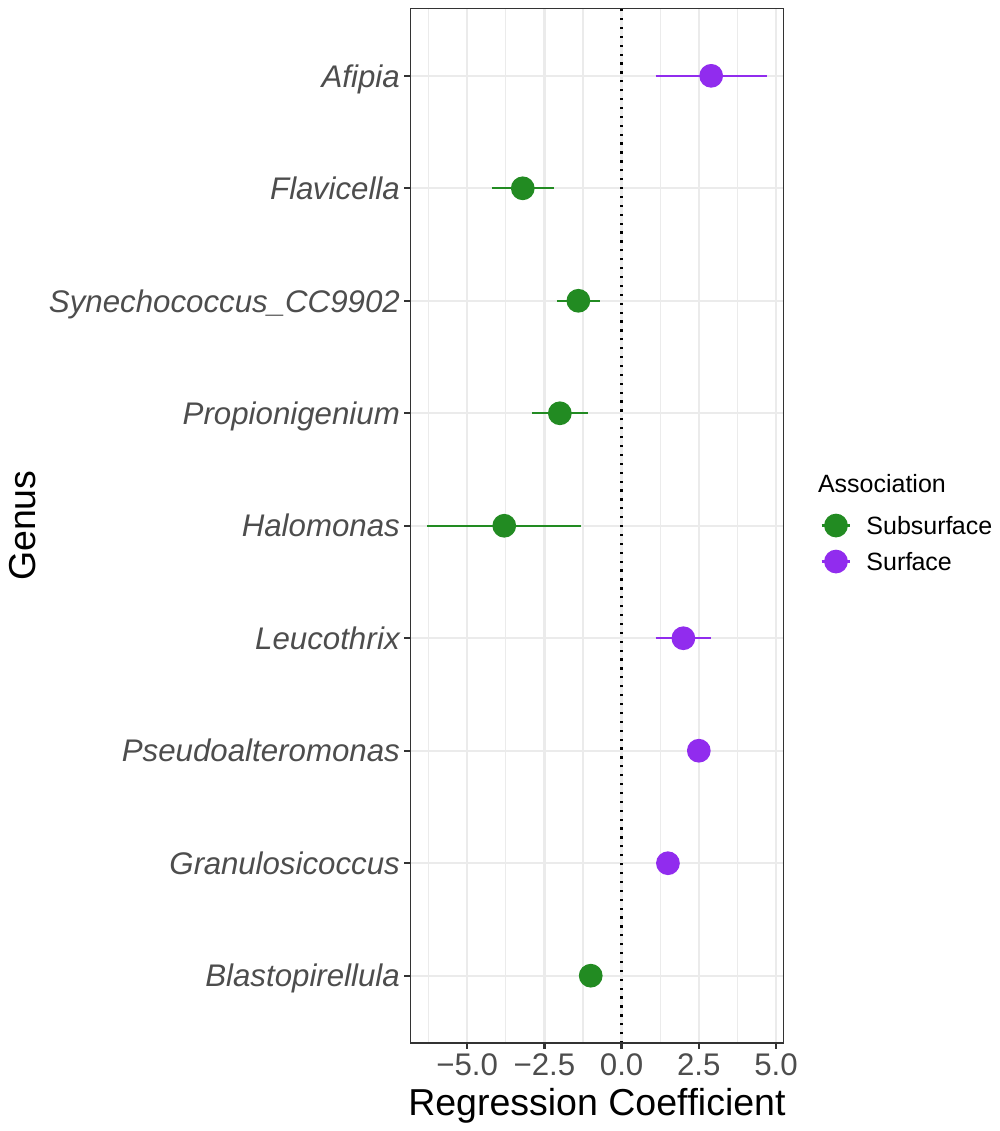

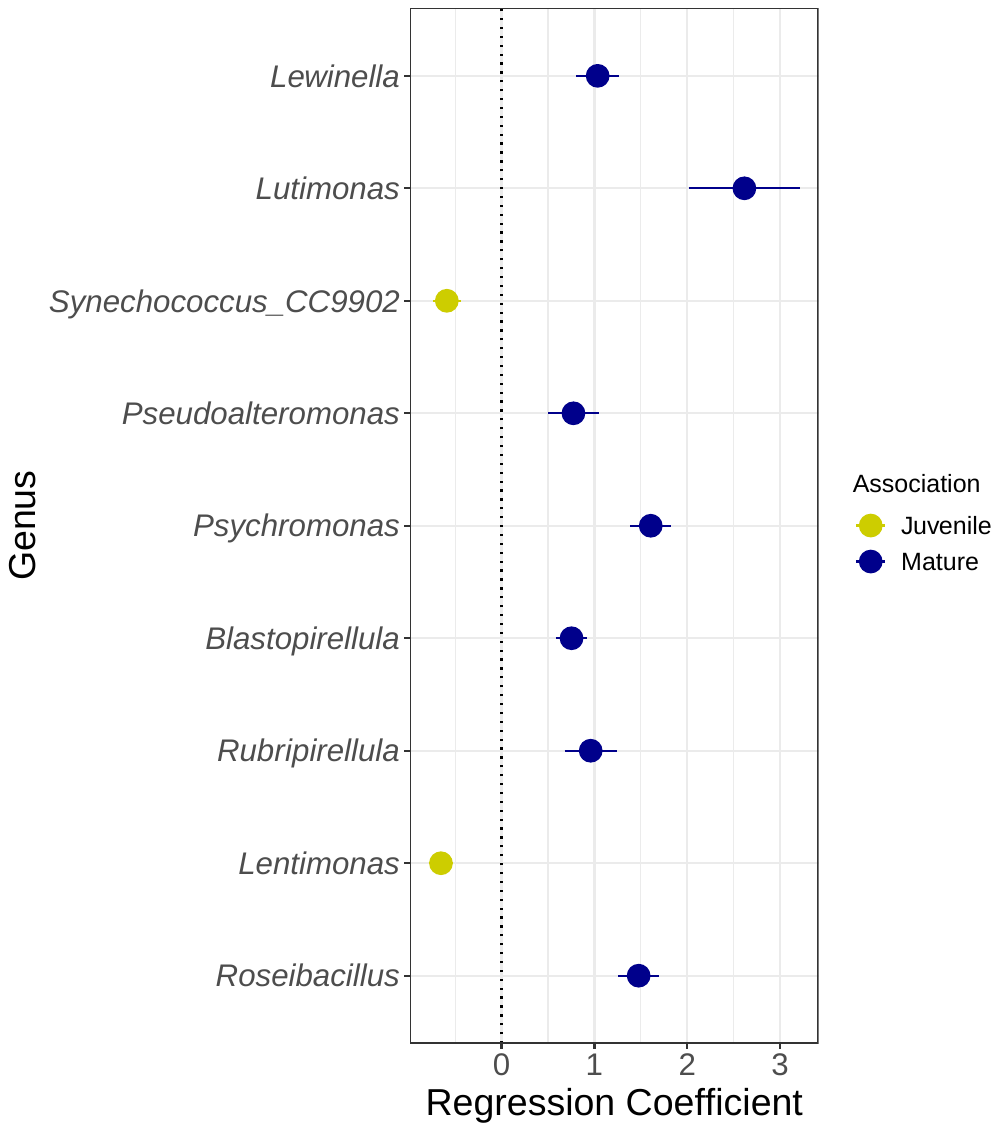

**SUPPLEMENTAL FIGURE 4:** Beta binomial regression analysis depicting differentially abundant bacterial genera associated with (A) age across all samples or (B) depth in mature samples exclusively. Among bacterial genera with relative abundance >0.5%, 9 are associated with age, and 9 are associated with depth. Genera are grouped alphabetically ordered by class.

B

**SUPPLEMENTAL FIGURE 5:** Beta binomial regression analysis depicting differentially abundant bacterial ASVs associated with (A) age across all samples or (B) depth in mature samples exclusively. Among bacterial ASVs with a prevalence ≥5 samples, 9 are associated with age, and 6 are associated with depth. ASVs are grouped alphabetically ordered by class.

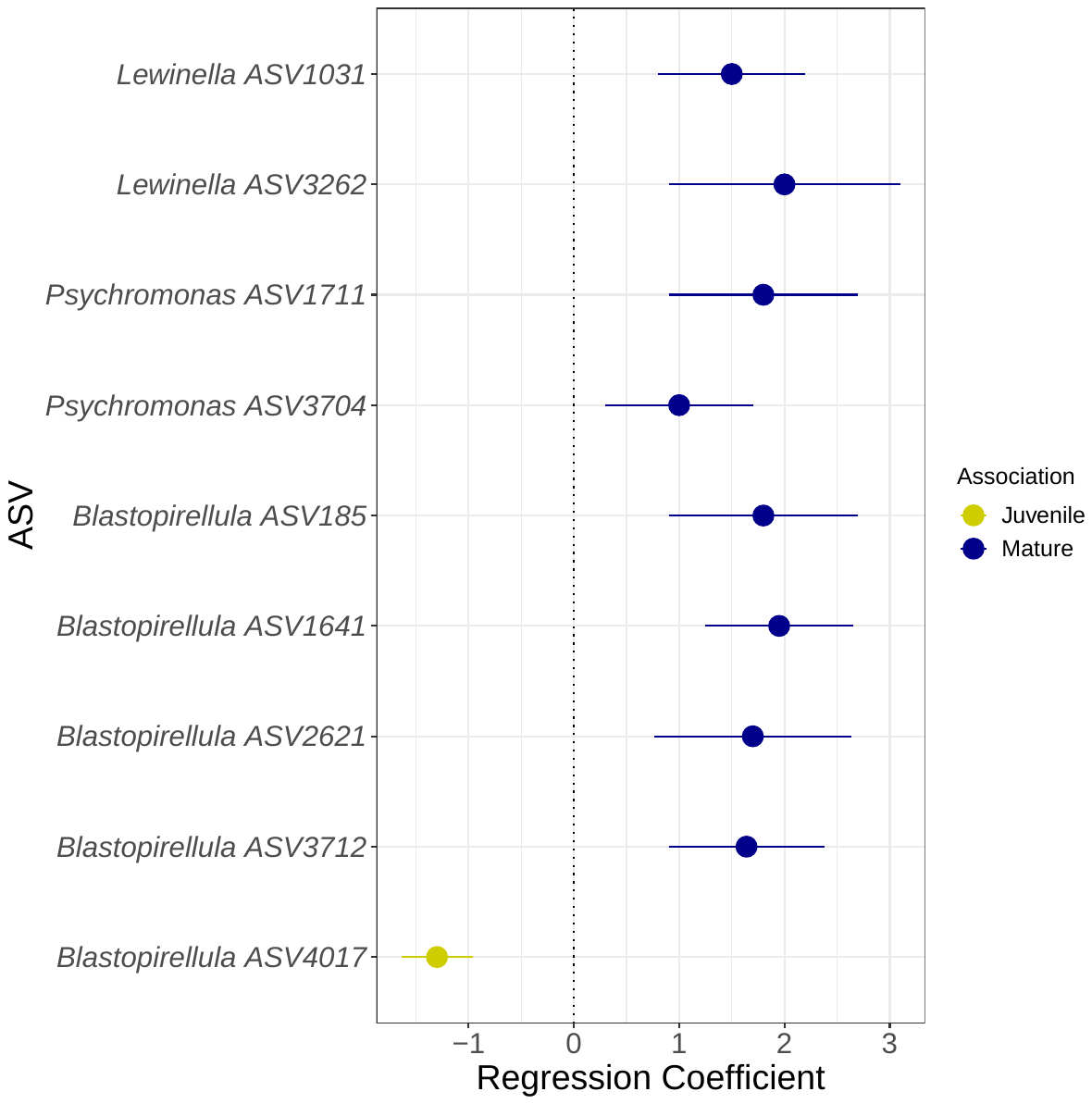

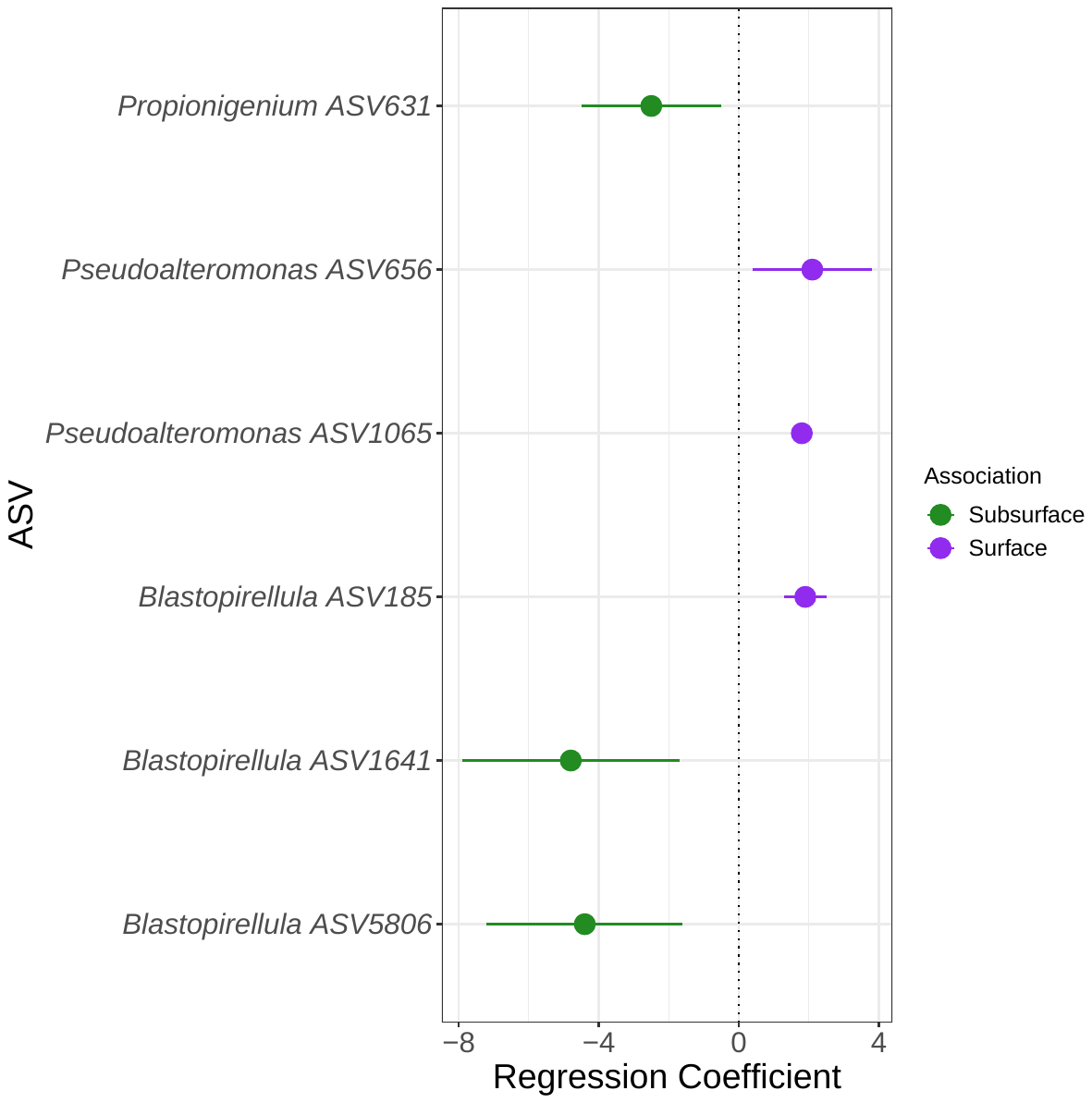

A

B

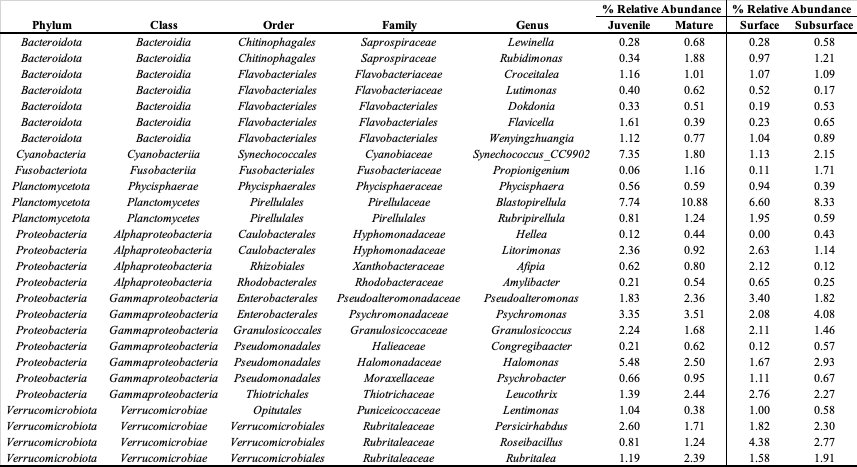

**SUPPLEMENTAL TABLE 3:** Percent relative abundances of all bacterial genera > 0.5% organized by depth and age categories.

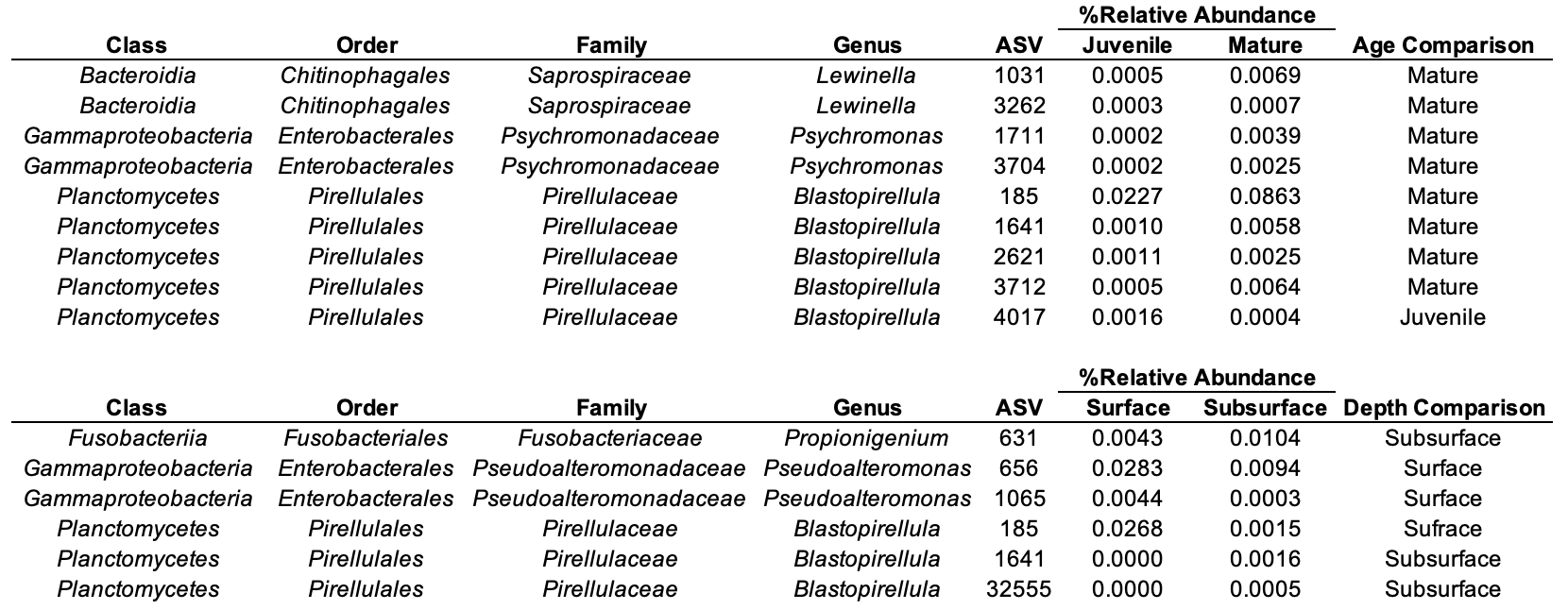

**SUPPLEMENTAL TABLE 4:** Percent relative abundances of differentially abundant bacterial ASVs identified by beta binomial regression (*P*<0.05). Columns display bacterial taxa, average percent relative abundances per category (i.e., age across all samples or depth in mature samples only), and indicate the category with significantly greater percent relative abundance.

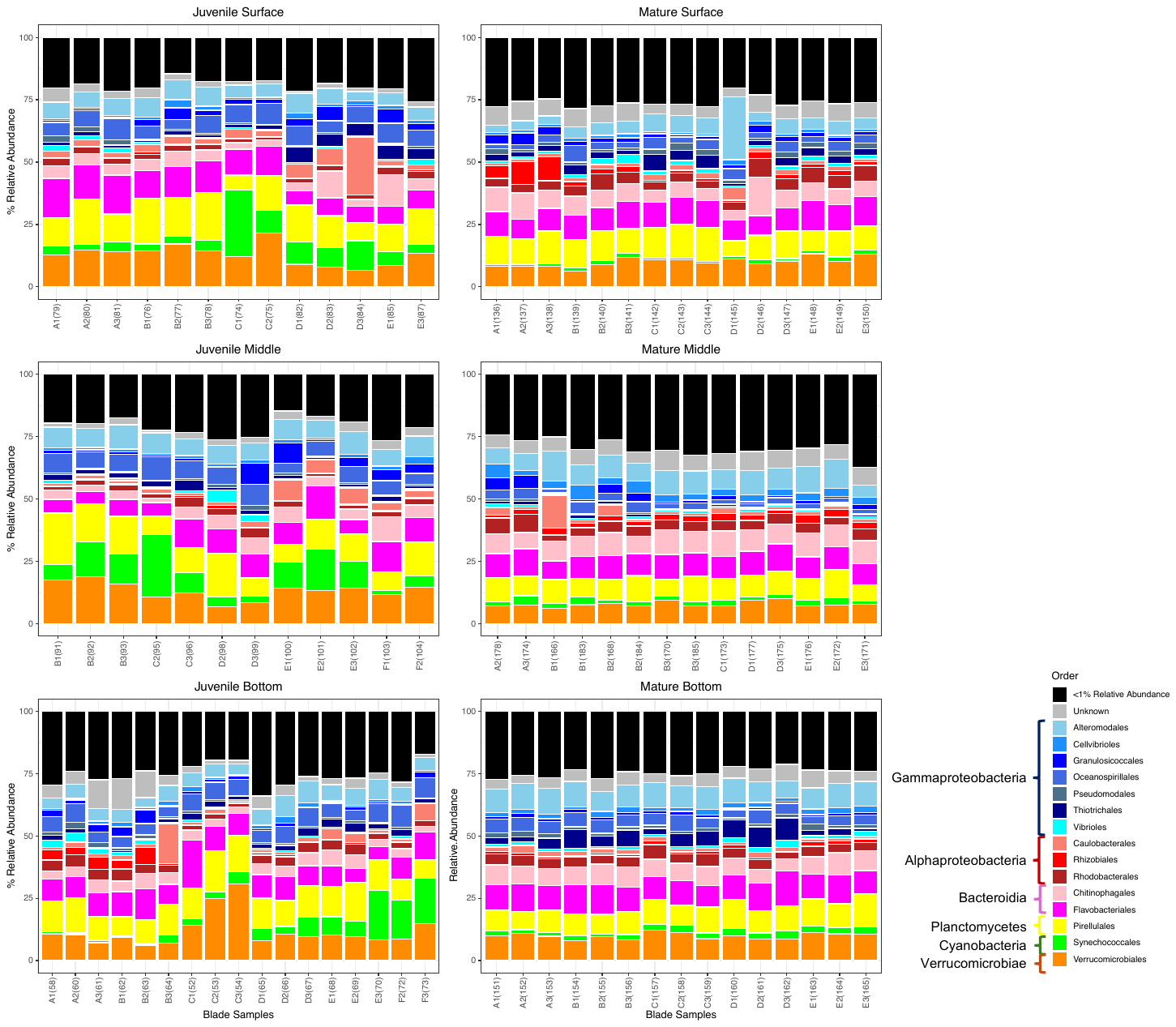

**SUPPLEMENTAL FIGURE 6:** Stacked bar plot of all bacterial orders with an average relative abundance > 1% across all samples. Each blade sample per depth and age category is represented as frond (e.g., A), blade number (e.g., 1), and unique sample id (e.g., 100). Black bars represent all orders with less than 1% relative abundance combined per sample. Gray bars represent all taxa with unknown orders. Bacterial orders are colored with similar shades per class.
